# Subject-specificity of the correlation between large-scale structural and functional connectivity

**DOI:** 10.1101/277590

**Authors:** J. Zimmermann, J. Griffiths, M. Schirner, P. Ritter, A.R. McIntosh

## Abstract

Structural connectivity (SC), the physical pathways connecting regions in the brain, and functional connectivity (FC), the temporal co-activations, are known to be tightly linked. However, the nature of this relationship is still not understood. In the present study, we examined this relation more closely in six separate human neuroimaging datasets with different acquisition and preprocessing methods. We show that using simple linear associations, the relation between an individual’s SC and FC is not subject-specific for five of the datasets. Subject-specificity of SC-FC fit is achieved only for one of the six datasets, the multi-modal Glasser HCP parcellated dataset. We show that subject-specificity of SC-FC correspondence is limited across datasets due to relatively small variability between subjects in SC compared to the larger variability in FC.

## Introduction

It has been shown that there is a relationship between structural connectivity (SC), the physical white-matter tracts between regions, and resting state functional connectivity (FC), the temporal coactivations between regions (Greicius, Supekar, Menon, & Dougherty, 2009; Hermundstad et al., 2013; Honey, Kotter, Breakspear, & Sporns, 2007; Honey et al., 2009; Koch, Norris, & Hund-Georgiadis, 2002; Misic et al., 2016; Ponce-Alvarez et al., 2015; Skudlarski et al., 2008; van den Heuvel, Mandl, Kahn, & Hulshoff Pol, 2009; van den Heuvel & Sporns, 2013) using both simple linear (Honey et al., 2009) as well as more complex metrics (Misic et al., 2016).

Most of this research, however, considers group-averaged matrices of SC and FC rather than individual connectomes. Motivated by the recent interest in personalized medicine and precision science, there is a greater need to understand individual differences and unique relationships between SC and FC. One important question is whether individual SC correlates with the corresponding subject’s FC to a greater extent than between-subjects. Correlations between whole-brain individual SC and FC have been associated with measures of behaviour or clinical conditions (Caeyenberghs, Leemans, Leunissen, Michiels, & Swinnen, 2013; Cocchi et al., 2014; Skudlarski et al., 2010; Zhang et al., 2011). Yet, there are very few studies that investigate the subject-specificity of this SC-FC correspondence (Honey et al., 2009; Meier et al., 2016), and as far as we know there are no studies that assert that individual SC maps best onto its corresponding FC using linear measures of association. One preliminary investigation conducted by Honey et al. (2009) examined this question, however, results were inconclusive due to the limited sample size. Clearly, it is not well understood whether there is a unique portion of variance in SC accounting for unique individual differences in FC.

It has already been shown that individual structural and functional connectomes can be sensitive to age (Zimmermann et al., 2016), personality traits (Markett et al., 2013), as well as cognition, demographics and behaviour (Hearne, Mattingley, & Cocchi, 2016; Ponsoda et al., 2017; S. Smith, 2016; S. M. Smith et al., 2015). Moreover, SC (Kumar, Desrosiers, Siddiqi, Colliot, & Toews, 2017; Munsell, 2017; Yeh et al., 2016) as well as FC (E. Amico, Goñi, J., 2017; Finn et al., 2015) can be used to identify individual connectome fingerprints. Nonetheless, the extent of this individual variability has been called into question (Marrelec, Messe, Giron, & Rudrauf, 2016; Waller et al., 2017), particularly for smaller sample sizes (Waller et al., 2017). For instance, it has been shown that variability in FC can be explained by only one or two dimensions, and that FC is highly degenerate in its ability to capture potential complexities and variability in underlying dynamics (Marrelec et al., 2016).

Variance decomposition methods, such as principal components analysis (PCA), are helpful for characterizing the strength of individual differences across connectomes (E. Amico, Goñi, J., 2017; Marrelec et al., 2016). PCA provides a simplified representation of the data by reducing the existing variance into a smaller number of components. In this way, the portion of variance that is common across subjects can be identified and separated from the unique aspects of variance.

The aim of the present study was to investigate the subject-specificity of the SC-FC relationship. The analyses were conducted on six datasets with variable acquisition schemes, preprocessing methods, and sample sizes (N = 48, 626, 171, 766, 754, 754). Four of these were variations of Human Connectome Project (HCP) data (Van Essen et al., 2013). We used simple linear measures of association with bootstrapping to quantify the correspondence of within-subject and between-subject SC-FC, and decomposition to quantify the extent of common and unique variability in SC and FC across subjects.

## Methods

### Data acquisition and preprocessing

The analyses were conducted on 6 MRI datasets of healthy subjects: the Berlin dataset (N = 48) (Ritter, Schirner, McIntosh, & Jirsa, 2013; Schirner, Rothmeier, Jirsa, McIntosh, & Ritter, 2015; Zimmermann et al., 2016), the Nathan Kline Institute (NKI) Rockland dataset from the UMCD Multimodal connectivity database (N = 171) (Brown, Rudie, Bandrowski, Van Horn, & Bookheimer, 2012), and four variations from the Human Connectome Project dataset (HCP) (S900 release) (Van Essen et al., 2013) that differed in terms of processing methods as well as parcellation schemes. These were: the HCP Lausanne dataset (N = 626), HCP Glasser dataset (N = 766), HCP Destrieux dataset (N = 754), and HCP Desikan-Killiany (DK) dataset (N = 754). Note that sample size differences between HCP datasets were due to removal of subjects with problematic parcellations. The HCP Glasser dataset was parcellated via a high-resolution multi-modal scheme based on an areal feature-based cross-subjects alignment method (Glasser et al., 2016). The research was performed in compliance with the Code of Ethics of the World Medical Association (Declaration of Helsinki). Written informed consent was provided by all subjects with an understanding of the study prior to data collection, and was approved by the local ethics committee in accordance with the institutional guidelines at Charité Hospital, Berlin, UCLA, and HCP WU-Minn.

A detailed description of data acquisition procedures is presented in Table S1. Subject sample size, age range, processing, and parcellation information are presented in Table 1, along with references to previously published papers with these datasets. Quality control was described in detail in those papers. For the Berlin and NKI Rockland dataset, noise-correction was performed via nuisance variable regression from the BOLD signal, including 6 motion parameters, mean white matter, and CSF signals. For the HCP datasets, we used FIX-denoised data, a tool that was trained to effectively remove components of the white matter, CSF, physiological noise, and 24 high-pass filtered motion parameters from the signal (Glasser et al., 2013).

**Table 1.** Dataset details for 6 MRI diffusion and resting-state datasets, including sample sizes, preprocessing methods, and parcellation schemes.

|  | **Berlin** | **NKI**<br>**Rockland** | **HCP,**<br>**Lausanne** | **HCP, Glasser** | **HCP,**<br>**Destrieux** | **HCP, DK** |
| --- | --- | --- | --- | --- | --- | --- |
| **Processing**<br>**reference** | Schirner et al. 2015 | Brown et al. 2012 | Glasser et al. 2013 | Glasser et al. 2013 | Glasser et al. 2013 | Glasser et al. 2013 |
| **Sample size** | 48 | 171 | 626 | 766 | 754 | 754 |
| **Subject ages** | 18-80<br>M = 41.90,<br>SD = 18.47) | 5-85<br>(M = 35.80,<br>SD = 19.99) | 22-36<br>(M = 28.65,<br>SD = 3.66) | 22-37<br>(M = 28.78,<br>SD = 3.70) | 22-37<br>(M = 28.78,<br>SD = 3.70) | 22-37<br>(M = 28.78,<br>SD = 3.70) |
| **Parcellation**<br>**(ROIs)** | Desikan-Killiany (68)<br>(Desikan et al., 2006) | Craddock (188) | Lausanne (83)<br>(Daducci et al., 2012) | Glasser (378)<br>(Glasser et al., 2016) | Destrieux (164)<br>(Destrieux, Fischl, Dale, & Halgren, 2010) | Desikan-Killiany (84)<br>(Desikan et al., 2006) |
| **Structural & diffusion processing** |  |  |  |  |  |  |
| **Software**<br>**method** | FreeSurfer | FreeSurfer & Dipy | TrackVis<br>Diffusion<br>Toolkit | HCP pipeline<br>(Glasser et al., 2013) | HCP pipeline<br>(Glasser et al., 2013) | HCP pipeline<br>(Glasser et al., 2013) |
| **Motion/eddy**<br>**correction** | yes | yes | yes | yes | yes | yes |
| **Intensity**<br>**normalized** | yes | no | yes | yes | yes | yes |
| **Tractography** | Probabilistic (MRTrix) | Deterministic (FACT) | Deterministic (EuDX) | Probabilistic (MRTrix) | Probabilistic (MRTrix) | Probabilistic (MRTrix) |
| **SC Metric** | Voxel pairs connected w streamline, ROI volume corrected | Streamline count | Streamline count | Weighted streamline count (SIFT2) by cross-sectional area | Weighted streamline count (SIFT2) | Weighted streamline count (SIFT2) |
| **Functional processing** |  |  |  |  |  |  |
| **Software**<br>**method** | Schirner et al 2015 | fMRI FEAT | Glasser et al 2013 | Glasser et al 2013 | Glasser et al 2013 | Glasser et al 2013 |
| **Slice-timing** | no | yes | no | no | no | no |
| **Motion** | MCFLIRT | MCFLIRT | FIX denoise | 6 DOF FLIRT, | 6 DOF FLIRT, | 6 DOF FLIRT, |
| ***correction*** |  |  |  | FIX denoise | FIX denoise | FIX denoise |
| ***Nuisance regression*** | 6 motion, mean WM, CSF | 24 motion, mean WM, CSF, volume | no | no | no | no |
| ***Smoothing*** | no | FWHM 5mm Gaussian | no | Cortical surface, subcortical volume, FWHM 2 mm Gaussian | Cortical surface, subcortical volume, FWHM 2 mm Gaussian | Cortical surface, subcortical volume, FWHM 2 mm Gaussian |
| ***Intensity normalization*** | no | yes | no | yes | yes | yes |
| ***Temporal filtering*** | High-pass 100s | Band-pass 0.08-0.009Hz | no | no | no | no |
| ***Registration to standard space*** | no | MNI152 | no | MNI152 & surface-based multimodal (MSMAll, Robinson et al., 2014) |  |  |
| ***Motion scrubbing*** | no | yes | no | no | no | no |

SC and FC was derived via diffusion-weighted magnetic resonance imaging (dwMRI) and resting-state blood oxygen dependent functional magnetic resonance imaging (rsfMRI BOLD), respectively. Structural and functional data were parcellated into predefined regions of interest (ROIs) that varied in size across datasets (68-378 cortical regions). Fibre track estimation was performed on the diffusion data, and weight and distance SCs were computed by aggregating tractography-based estimations of white matter streamlines between ROIs. Each entry in the SC weights matrix was an estimate of the connection strength between a pair of ROIs. SC distances were the Euclidian distances (Brown et al., 2012; Glasser et al., 2013; Hagmann et al., 2008), or average length of tracks (Schirner et al., 2015) in mm between pairs of ROIs. We corrected for SC distance by regressing distances from weight SCs, and using residuals for analysis (as tract length may have an effect on structure-function relations (Romero-Garcia, Atienza, & Cantero, 2014)). To account for age-related differences in parcellation and ROI size in the Berlin dataset, SCs were weighted by the mean gray-matter white-matter interface area of connected ROIs. FCs were computed as the Pearson’s correlation between each ROI pair of BOLD time series, and were transformed to a normal distribution via a Fisher’s r to Z.

### Subject-specificity of SC-FC predictions

We compared individual SC and FC within and between all subjects using Pearson’s correlations, in order to determine whether individual SC correlates best with its own individual FC. We constructed a matrix of size N_SC_ x N_FC_ (N = the number of subjects, N_SC_ = N_FC_). The diagonal of this matrix captures the intra-subject (within) SC-FC correlations; the off-diagonal represents the inter-subject (between) SC-FC (See Figure 1 for a visualization of this SC-FC matrix). We corrected the p-value of each correlation value in the resulting matrix for multiple comparisons using FDR (Matlab function fdr_bky) (Benjamini, Krieger, & Yekutieli, 2006). Note that associations between all individual SC and FC within and between all subjects was also performed via eigenvector correlations, to complement the Pearson’s correlation method. This method is described in the Supplementary Materials.

**Figure 1.**
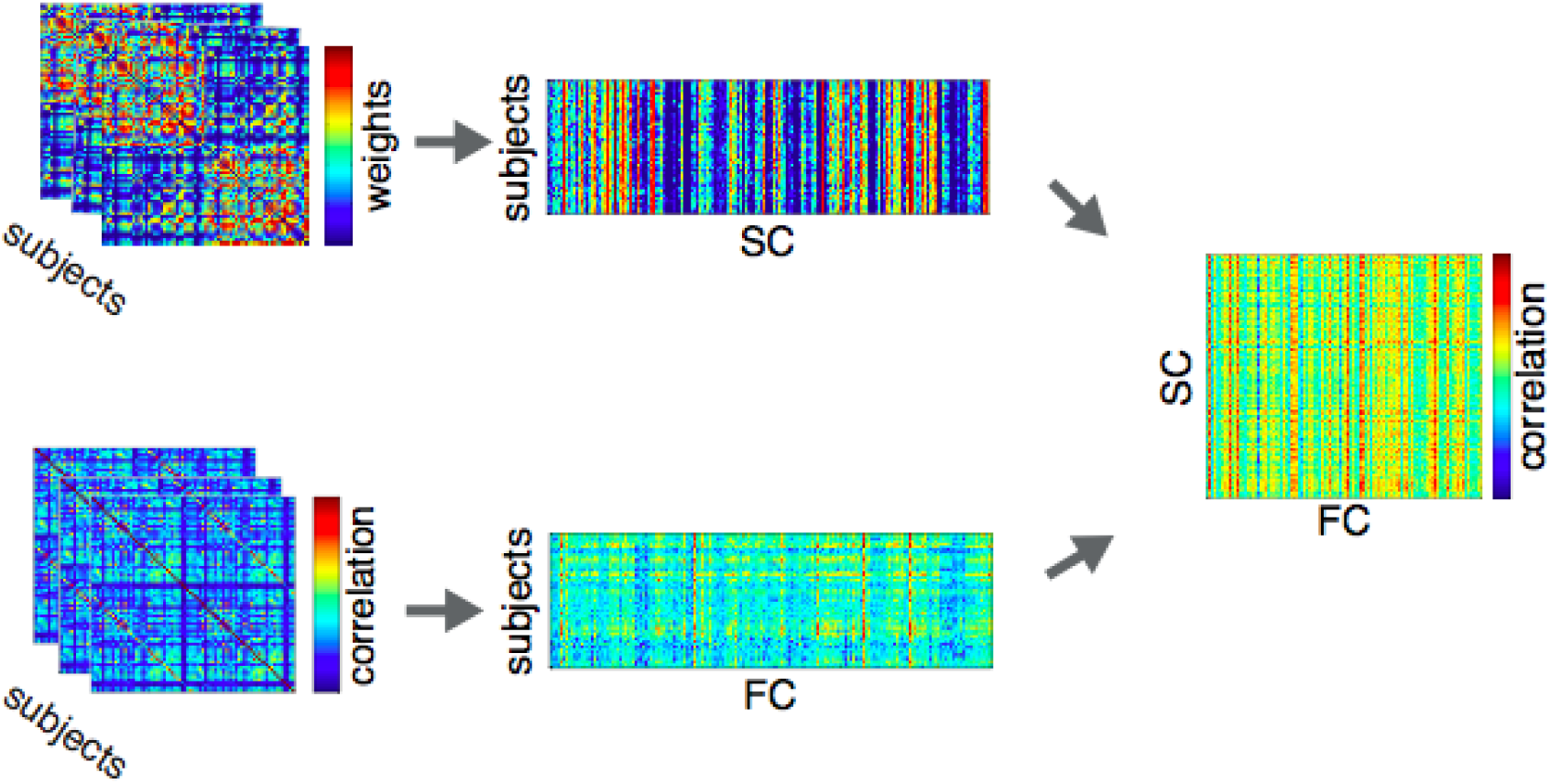
Individual subject SCs and FCs were stacked into a subjects x connections matrix. Subject-wise SC and FC were then correlated for all pairs of SC and FC (within and between subjects). On-diagonals of this matrix represent within-subject SC-FC (SC-SelfFC), off-diagonals represent between-subject SC-FC (SC-OtherFC).

We conducted 1000 bootstrapped means of SC-SelfFC correlations and 1000 bootstrapped means of SC-OtherFC correlations (Matlab function bootstrp), plotting the two bootstrapped distributions against each other. In order to evaluate the statistical significance of the differences between the distributions, we subtracted the SC-otherFC distribution from the SC-selfFC distribution and constructed a 95% confidence interval on this difference distribution (Matlab function prctile). To confirm our results, we conducted several secondary analysis. Firstly, we conducted a global signal regression in order to minimize the effects of global signal differences on individual SC-FC relationships. We also logarithmized SCs and transformed them to a Gaussian distribution by resampling (Honey et al., 2009) to correct for exponentially distributed connection weights. Lastly, we used only SC present connections, as indirect connections may have an unknown effect on FC (Honey et al., 2009).

### Subject variability in SC and in FC

We examined variability across subjects in SC as well as in FC via PCA. The objective was to understand whether the lack of subject-specificity of SC-FC in the Berlin, HCP Lausanne and NKI Rockland dataset was due to a large portion of common variance in the connectomes across subjects over-powering existing individual differences. To this end, we decomposed the subject-wise SC matrix (SC connections x subjects) and FC matrix (FC connections x subjects) via the Matlab princomp function (subjects as variables). The breakdown of variability in SC as well as FC across subjects was thus ascertained. From the PCA, we obtained for each PC: eigenvalues, principal component loadings per subject, and principal component scores per each connection. To determine the significance of the resulting eigenvalues, we generated null distributions of eigenvalues for each PC by permuting the SC and the FC 100 times (scrambled across connections and subjects) and performing PCA of the resulting matrices. A p-value for each PC eigenvalue was obtained as the proportion of times that the permuted eigenvalue exceeded the obtained eigenvalue.

We also computed the age effect on connectome variability by calculating the correlation, via Partial Least Squares, of age (age vector, size: subjects x 1) with the subjects’ principal coefficient loadings of the significant PCs (size: subjects x number of significant PCs). Partial Least Squares is a multivariate method comparable to canonical correlation in that it computes the relationship between two matrices via orthogonal latent variables (Krishnan, Williams, McIntosh, & Abdi, 2011; McIntosh & Lobaugh, 2004). The significance of the resulting correlations was assessed via permutation testing (N = 1000) of the singular values from singular value decomposition of the two matrices. Reliability of each principal component subject loading to the latent variable was assessed via bootstrapping (N = 500). We thus were able to compute how age corresponded to the significant variance across subjects.

## Results

### Subject-specificity of SC-FC predictions

We first quantified the SC-FC relationship at the group-average level. The correlation between averaged SC and averaged FC was as follows: *r* = 0.59, 0.47, 0.41, 0.34, 0.40, 0.47, *p* < 0.001, for Berlin, HCP Lausanne, NKI, HCP Glasser, HCP Destrieux, and HCP DK, respectively. At the individual subject level, all subjects’ SCs were significantly correlated with all subjects’ FCs (between and within SC-FC) (Pearson’s correlations*, p* < 0.001, FDR multiple comparison correction, *p* < 0.001). However, we found that SC-FC correlations were subject-specific only for the HCP Glasser dataset, and not for the other datasets. This was assessed by comparing the bootstrapped within-subject SC-FC correlation distribution (SC-SelfFC) with the bootstrapped between-subject SC-FC correlation distribution (SC-OtherFC), as discussed in the Methods. See Figure 2 for subject-specificity assessed using the simple bivariate correlation, and Figure S2 for subject-specificity assessed using the eigenvector correlation approach. Mean and CIs on the difference distributions are shown in Table 2 below for simple correlations and Table S2 for eigenvector correlations. The results were consistent using the two approaches.

**Table 2.** Mean and 95% CIs on the difference distribution, calculated as the difference between the SC-SelfFC and SC-OtherFC distributions. The * indicates a significant subject-specificity, whereby the distribution of intra-subject SC-FC was higher than the distribution of inter-subject SC-FC.

| Dataset | Simple correlation |  |
| --- | --- | --- |
|  | Mean | CI |
| Berlin | M = 0.0013 | [-0.0169, 0.0190] |
| HCP, Lausanne | M = 0.0016 | [-0.0012, 0.0044] |
| NKI Rockland | M = -5.8447e-04 | [-0.0109, 0.0065] |
| HCP, Glasser | M = 0.0032 | [0.002, 0.0043] * |
| HCP, Destrieux | M = -2.2348e-04 | [-0.0019, 0.002] |
| HCP, DK | M = 0.001 | [-.0001, 0.0017] |

**Figure 2.**
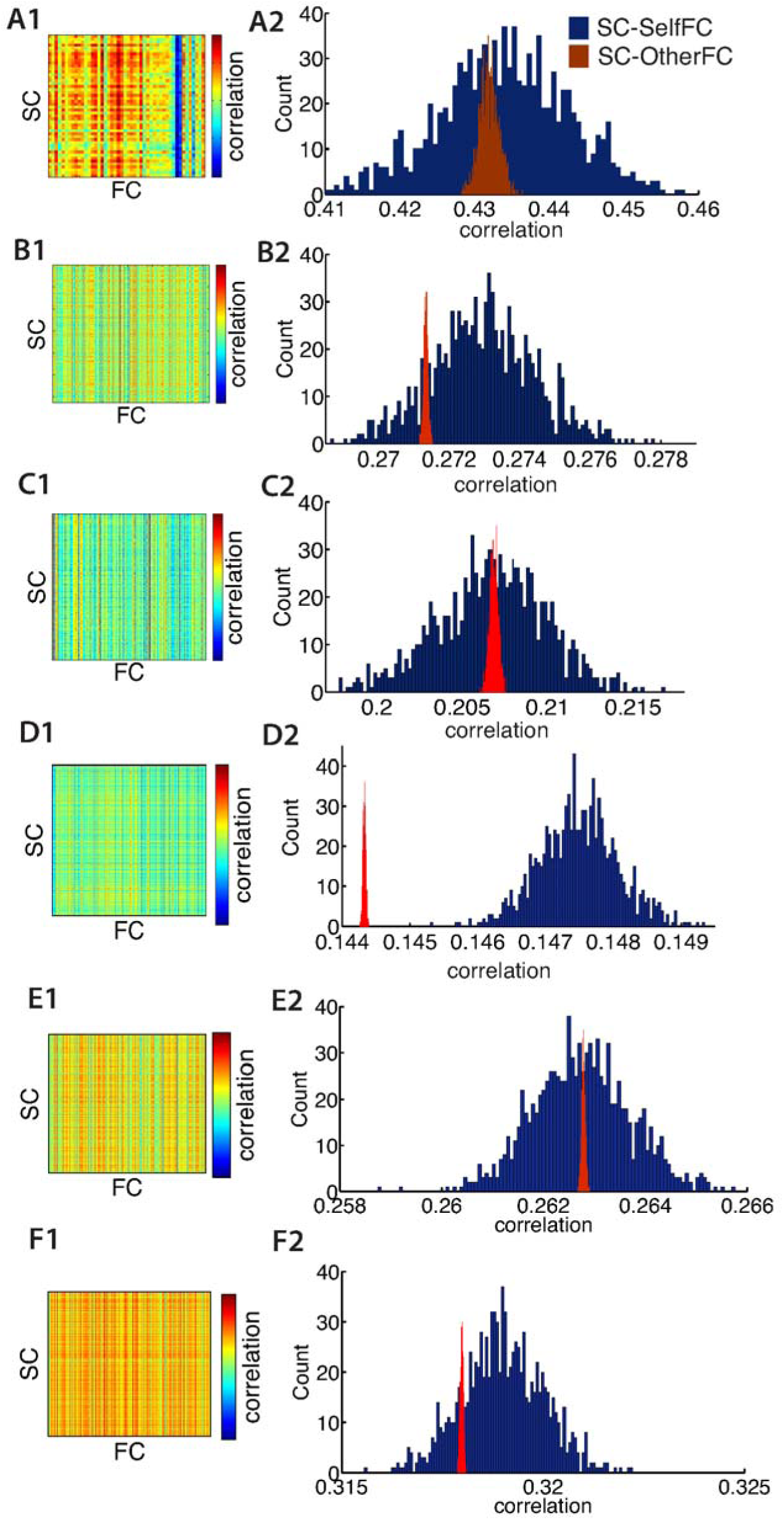
Bivariate Pearson’s correlations are shown for all combinations of SC and FC within and between subjects in the first column. Distribution histograms of bootstrapped means of intra (SC-SelfFC) and inter (SC-OtherFC) correlations are shown in the second column. Each row is a different dataset: **A)** Berlin **B)** HCP Lausanne **C)** Rockland **D)** HCP Glasser, **E)** HCP Destrieux **F)** HCP DK.

In summary, we found that for all but the HCP Glasser dataset, a subject’s SC did not correlate better with its own FC than with another subject’s FC. These results remained consistent when using distance corrected SCs, or only SC present connections. For the HCP Glasser dataset, the within-subject SC-FC was significantly higher than the between-subject SC-FC.

### Subject variability in SC and in FC

Figures 3 through 8 show PCA results for Berlin, HCP Lausanne, NKI, HCP Glasser, HCP Destrieux, and HCP DK data, respectively. For both SC and FC across our datasets, the first component captured a very large portion of common variance across subject. All subjects loaded heavily on this common PC1; these principal component subject loadings are visualized in the bar plots on the right-hand-side of Panel B in Figures 3-8. The principal component scores (i.e., reconstructed matrix from PC1) for this common PC are visualized in the matrices on the left-hand-side of Panel B in Figures 3-8. These represent the features of the connectome that were captured by PC1. The variance explained by this first common PC was large in the SC (91%, 80%, 79%, 91%, 93% variance explained for Berlin, HCP Lausanne, NKI, HCP Glasser, HCP Destrieux, and HCP DK datasets respectively) and lower in the FC (57%, 70%, 33%, 74%, 80% variance explained for Berlin, HCP Lausanne, NKI, HCP Glasser, HCP Destrieux, and HCP DK datasets respectively). Eigenvalues for the first 30 PCs for all datasets are shown in Supplementary Table S3. It is noteworthy that the HCP Glasser SC showed the largest *number* of significant principal components (HCP Glasser N = 12, Berlin = 1, HCP Lausanne = 7, NKI Rockland = 2, HCP Destrieux = 8, HCP DK = 7).

**Figure 3.**
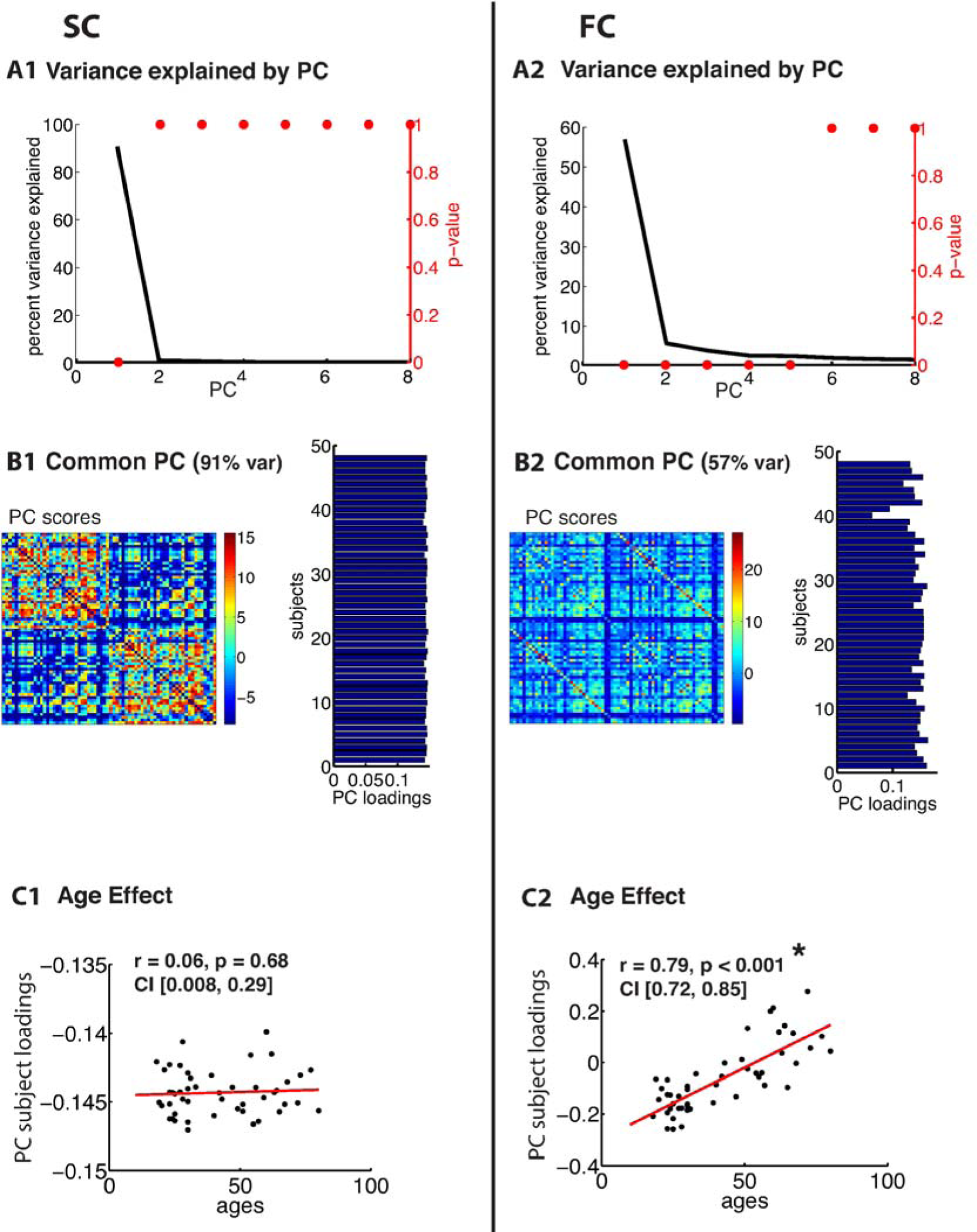
PCA of SC (left column) and FC (right column) for the Berlin dataset. **A)** The first row depicts the percent of total variance explained for each principal component (PC) with corresponding p-values in red. **B)** The second row shows the PC connectome scores as well as the individual subject loadings on the first PC (all subjects positively loaded). **C)** The third row shows the effect of age.

**Figure 4.**
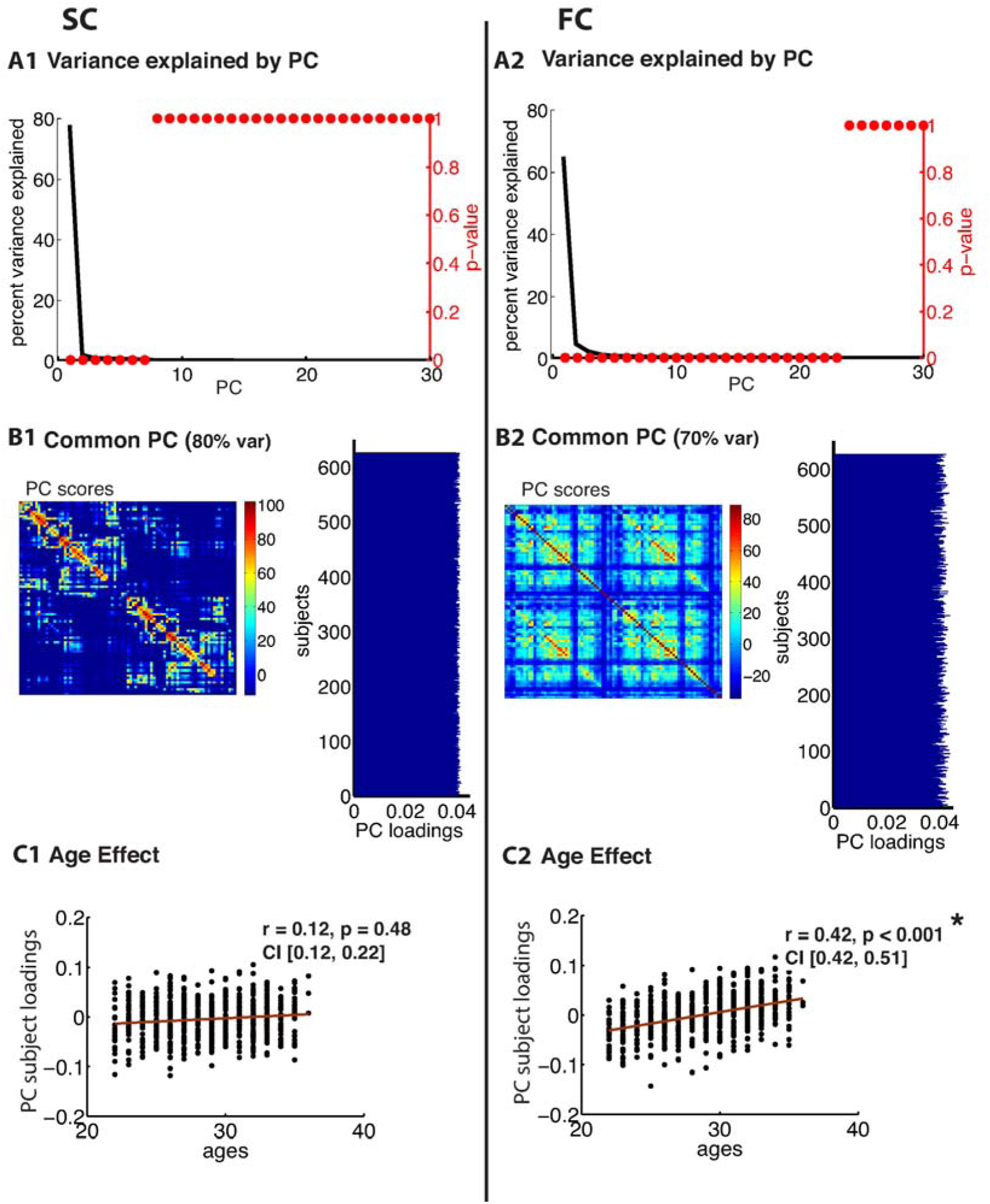
PCA of SC (left column) and FC (right column) for the HCP Lausanne dataset. **A)** The first row depicts the percent of total variance explained for each principal component (PC) with corresponding p-values in red. **B)** The second row shows the PC connectome scores as well as the individual subject loadings on the first PC (all subjects positively loaded). **C)** The third row shows the effect of age.

**Figure 5.**
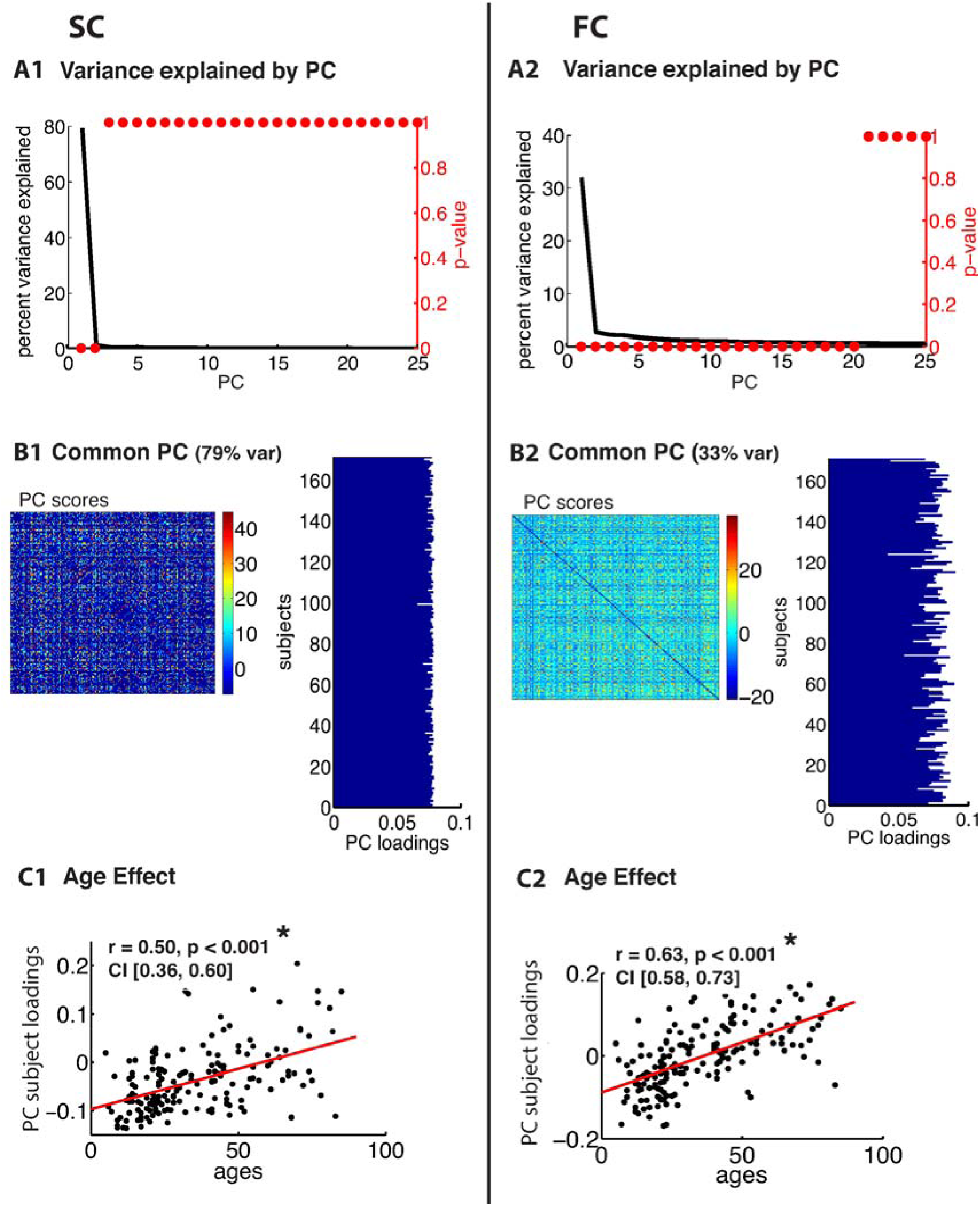
PCA of SC (left column) and FC (right column) for the Rockland dataset. **A)** The first row depicts the percent of total variance explained for each principal component (PC) with corresponding p-values in red. **B)** The second row shows the PC connectome scores as well as the individual subject loadings on the first PC (all subjects positively loaded). **C)** The third row shows the effect of age.

**Figure 6.**
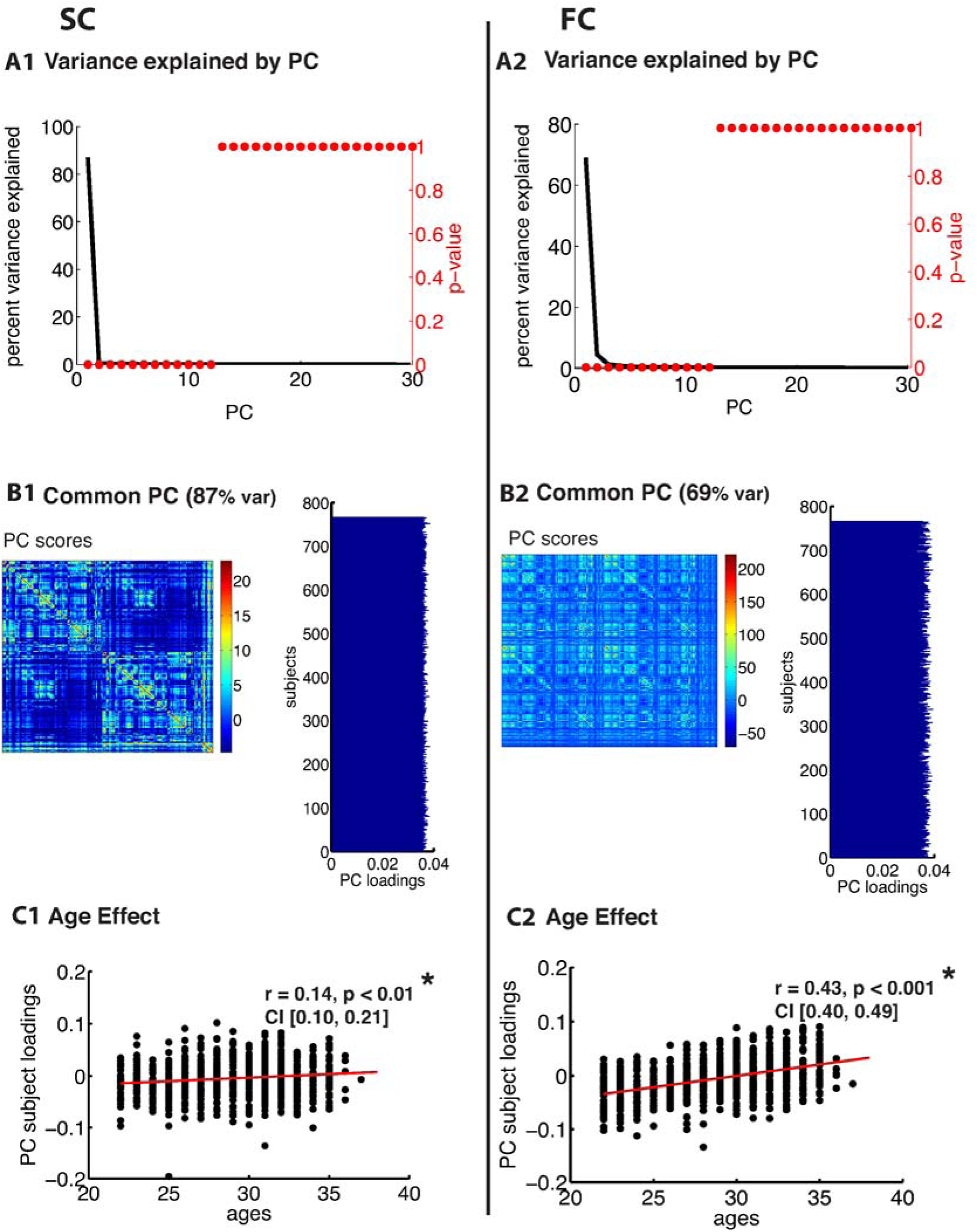
PCA of SC (left column) and FC (right column) for the HCP Glasser dataset. **A)** The first row depicts the percent of total variance explained for each principal component (PC) with corresponding p-values in red. **B)** The second row shows the PC connectome scores as well as the individual subject loadings on the first PC (all subjects positively loaded). **C)** The third row shows the effect of age.

**Figure 7.**
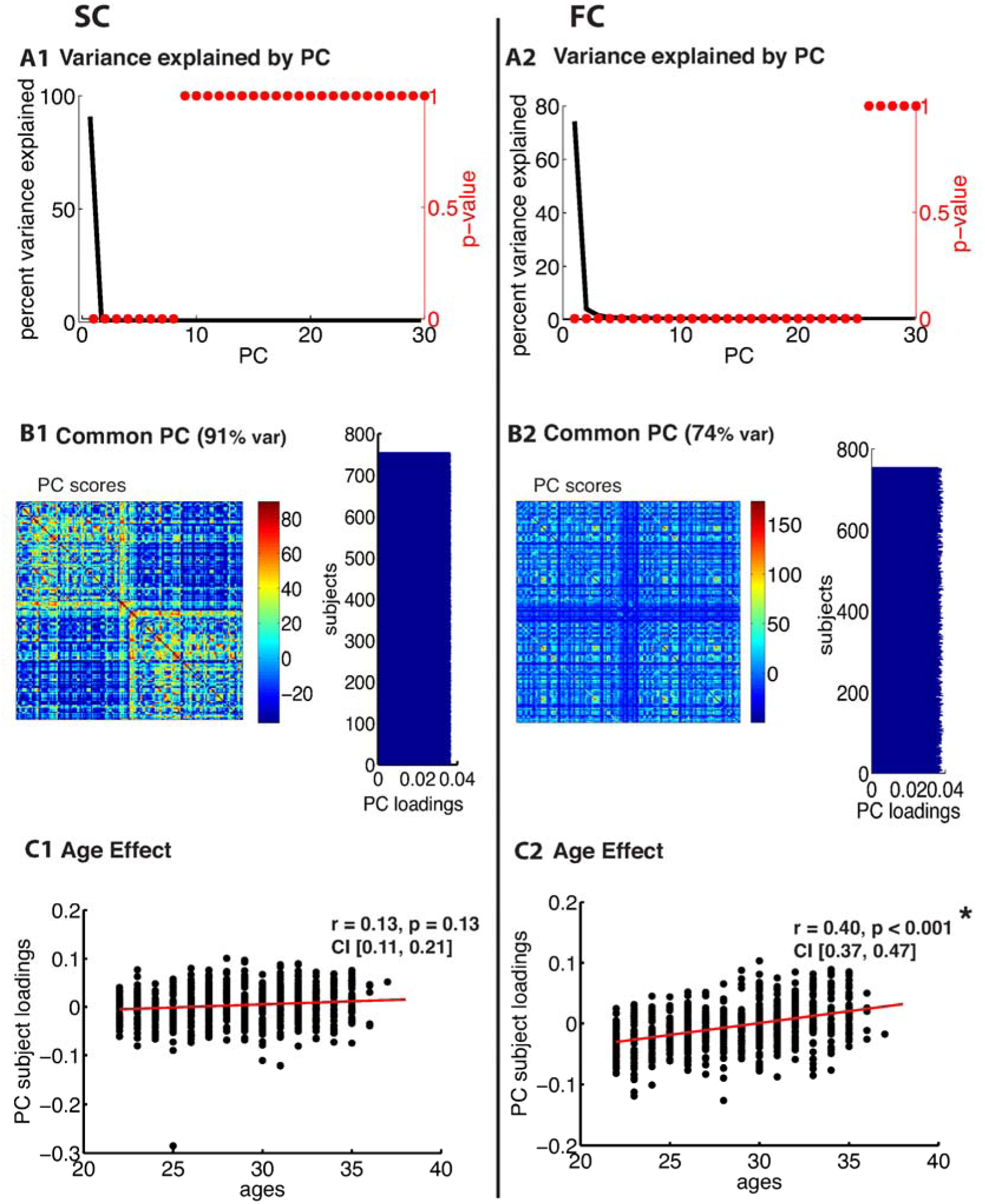
PCA of SC (left column) and FC (right column) for the HCP Destrieux dataset. **A)** The first row depicts the percent of total variance explained for each principal component (PC) with corresponding p-values in red. **B)** The second row shows the PC connectome scores as well as the individual subject loadings on the first PC (all subjects positively loaded). **C)** The third row shows the effect of age.

**Figure 8.**
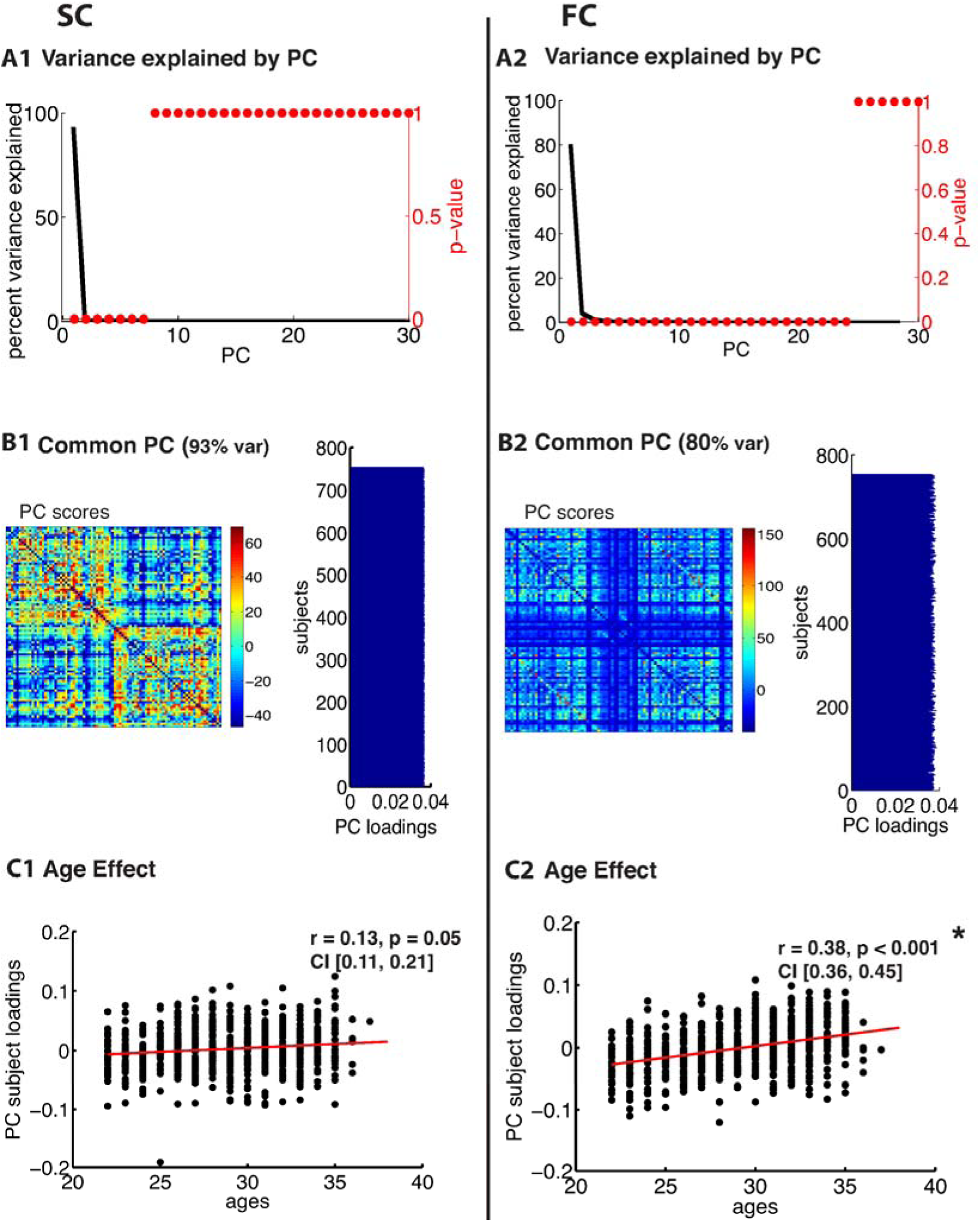
PCA of SC (left column) and FC (right column) for the HCP DK dataset. **A)** The first row depicts the percent of total variance explained for each principal component (PC) with corresponding p-values in red. **B)** The second row shows the PC connectome scores as well as the individual subject loadings on the first PC (all subjects positively loaded). **C)** The third row shows the effect of age.

A second pattern of results that we observed across all datasets was that SC was less variable than FC across subjects. There were fewer significant eigenvalues for the SC compared to the FC (See Table S3). From the figures (Panel A in Figures 3-8), the knee, or drop-off in the variance explained by subsequent PCs (Cattell, 1966) was evidently sharper for the SC than the FC. Thus, although the common component was dominant for both modalities, the second and later components explained a larger portion of variance in the FC than in the SC.

Consistent with the above findings, we also noted differences between SC and FC in the strength of the age-related differences. We found an age effect in the FC for all 6 datasets (Berlin: *r* = 0.79, *p* < 0.001, HCP Lausanne: *r* = 0.42, *p* < 0.001, NKI: *r* = 0.63, *p* < 0.001, HCP Glasser: *r* = 0.43, *p* < 0.001, HCP Destrieux: *r* = 0.40, *p* < 0.001, HCP DK: *r* = 0.38, *p* < 0.001). We found an age effect in the SC for 2 of the 6 datasets (Berlin (non-significant): *r* = 0.06, *p* = 0.68, HCP Lausanne (non-significant): *r* = 0.12, *p* = 0.48, NKI (significant): *r* = 0.50, *p* < 0.001, HCP Glasser (significant): *r* = 0.14, *p* = 0.035, HCP Destrieux (non-significant): *r* = 0.13, *p* = 0.13, and HCP DK (non-significant): *r* = 0.13, *p* = 0.05).

We compared brain volume across subjects to check for any age-related differences. For the Berlin and the Rockland dataset, tissue segmentation was performed and partial volume maps were derived using FSL FAST. Total brain volume was computed by summing the GM and WM tissue volumes. Total brain volume across subjects was correlated with region-wise SC (Berlin: *r*= 0.22, *p* = 0.14, Rockland: *r* = 0.18, *p* = 0.17) and FC (Berlin: *r* = 0.17, *p* = 0.31, Rockland: *r* = 0.09, *p* = 0.40); no effect was found. Volume differences in the HCP data were already accounted for via the FIX method.

Finally, it is noteworthy that our results remained robust following a number of secondary analyses. These are described in detail in the Methods, and include the following: global signal regression, including only SC present connections in the analyses, and logarithmizing and resampling SCs to a Gaussian distribution. Because results remained robust against these corrections, the results shown are those based on the original matrices. Please see Supplementary Table S2 for the PCA results on logarithmized SCs redistributed to Gaussian.

## Discussion

### Subject-specificity in SC-FC

Initial studies of SC-FC correspondence (Greicius et al., 2009; Honey et al., 2007; Honey et al., 2009; Koch et al., 2002) show that there is a relationship between these two entities via linear (Honey et al., 2009) as well as more complex methods (Misic et al., 2016). However, there remains a gap in our understanding of how the two measures are related at the individual level. In the present study, we showcase how individual SC corresponds with individual FC using simple linear metrics in six separate datasets (Berlin, HCP Lausanne, NKI Rockland, HCP Glasser, HCP Destrieux, HCP DK). The datasets differed in sample size, acquisition and processing methods as well as age spectrums. The question was whether the correspondence of individual SC-FC matrices was greater than if two matrices were randomly paired.

Our results showed that, although there is a correlation between group-averaged SC and FC, replicating previous findings (Greicius et al., 2009; Hermundstad et al., 2013; Honey et al., 2007; Honey et al., 2009; Koch et al., 2002; Misic et al., 2016; Ponce-Alvarez et al., 2015; Skudlarski et al., 2008; van den Heuvel et al., 2009), the specificity of this SC-FC relationship was not unique to an individual. Five of the datasets examined did not show subject-specificity of the SC-FC correspondence, so that within-subject SC-FC did not exceed random pairings of SC-FC. This would suggest that individual SC cannot predict individual FC beyond chance. However, when the analysis was conducted on the HCP data with the Glasser parcellation, significant subject-specificity was observed. This would suggest that while subject-specificity assessed on standard datasets via standard parcellation and processing methods is difficult to ascertain, it may be obvious only when higher resolution data *as well as* finer parcellations are used.

Our finding that intra-subject SC-FC correspondence exceeded inter-subject SC-FC correspondence for the HCP Glasser dataset, but not for the remaining datasets, supports the hypothesis by Honey et al. (2009). Honey et al. (2009) speculated that the individual SC-FC fit would be significant if shown on a large enough dataset of high fidelity. However, fidelity of the data will depend on a number of factors, including the quality and rigor of the data acquisition procedures, the processing methods (e.g. tractography), and the parcellation used. The acquisition procedures alone were unlikely to be the sole driving factor behind subject-specificity, as these were consistent across the HCP data. We hypothesized that the superior subject-specificity of the Glasser HCP data (compared to the HCP Lausanne) was due to the high-precision parcellation used (Glasser et al., 2016). However, these two HCP datasets also differed in the tractography method (probabilistic versus deterministic). Thus we endeavoured to re-evaluate our findings post-hoc using two additional HCP datasets with probabilistic tractography processed in the same way as the Glasser HCP, except with different parcellation methods. We used the FreeSurfer convolution-based probabilistic Destrieux atlas (Destrieux et al., 2010) and the Desikan-Killiany (DK) atlas (Desikan et al., 2006). We did not find subject-specificity with the HCP Destrieux and the HCP DK, suggesting that the Glasser parcellation allows for a fitting of individual structure and function that could not otherwise be observed. The Glasser multi-modal parcellation is based on functional properties with improved areal feature-based cross-subject alignment, rather than solely geometric and morphological properties. Thus, the method improves the neuroanatomical precision of individual parcellations. It is important to point out that despite the improvement, the HCP Glasser dataset was only slightly better than the others, and would not pass a direct head-to-head comparison since the presence of significance in one dataset and the absence of significance in another does not mean the two datasets are themselves significantly different.

### Subject-specificity in SC-FC is limited by variability within modality

The second set of findings showed that the unique portion of variance that exists in either modality alone is limited. This may restrict the portion of SC that can reasonably be captured by individual FC. We had hypothesized that the lack of subject-specificity in the Berlin, HCP Lausanne, NKI Rockland, HCP Destrieux, and HCP DK dataset was due to a large portion of common variance in the connectomes across subjects that over-powered any existing individual differences. Indeed, our results conferred that there is a large portion of common variance in SC across subjects. This was the case regardless of the sample size, data quality, or parcellation. Interestingly, even in the Glasser dataset, where SC-FC subject-specificity was observed, the common component was strikingly large. We did observe, however, that SC variability was captured via a greater *number* of components in the Glasser dataset compared to the other datasets, suggesting greater inter-individual differences in the SC. Although the smaller datasets (e.g. Berlin) generally had fewer SC components, the variability that was observed in the HCP Glasser SC was not merely due to sample size, as the HCP Destrieux and HCP DK datasets were comparable to the HCP Glasser dataset in terms of sample size.

We also observed a large common component in the FC across subjects. However, this component accounted for a smaller portion of total variance than the SC common component. Moreover, a smaller *number* of significant variance components were found in the FC across subjects compared to in the SC. Together, these results suggest that FC is more variable than SC across subjects. This can also be observed in the striping of the SC-FC correspondence matrix, where some FCs correlate strongly with all SCs, while others correlate very little with all SCs. Note that this does not mean that individual differences in SC were not observed, but rather that the variance in SC that maps onto the corresponding variance in FC is weaker than one may expect intuitively.

In the FC, a significant portion of variance was related to age, particularly for the two datasets with a wide age range (Berlin, NKI: age = 20-80, 5-85). This is consistent with previous reports of age effects on FC (Andrews-Hanna et al., 2007; Damoiseaux et al., 2007; Ferreira & Busatto, 2013; Sala-Llonch et al., 2014). Interestingly, age did not account for a significant portion of between-subjects variance in SC for four of the six datasets. We found an age effect in the SC only for the HCP Glasser dataset and the NKI Rockland dataset. In the NKI Rockland dataset, the large observed age effect in SC was likely a consequence of the wide age distribution and the inclusion of child subjects. The grey-white matter boundary is ill defined in children, and incomplete myelination results in weaker tractography-based estimation of SC (Deoni, Dean, Remer, Dirks, & O’Muircheartaigh, 2015; Thompson et al., 2005).

The limited amount of between-subjects variability in both SC and FC that we observed was comparable to findings by Marellec et al. (2016), where a large portion of variance was accounted for by an invariant core that was consistent across subjects (SC: ~86%, FC: ~59%). There, it was shown that the invariant core of SC correlated with the invariant core of FC. Along the same lines, Waller et al. (2017) suggested that the specificity of connectome fingerprinting using FC was limited by the large amount of common variance across subjects.

The decomposition approach we used here was helpful for separating common and unique variance, and identifying aspects of the connectome that express each portion of variance. Data-driven classification algorithms like clustering are an alternate approach that can be used to express similarities and differences between subject connectomes (E. Amico et al., 2017; Iraji et al., 2016). Recently, a consensus clustering algorithm has been introduced that can be helpful for identifying how aspects of the connectome are combined to express these inter-subject similarities and differences (Rasero, 2017).

### Limitations on the study of variability within modality

The study of variability within SC and FC each faces its unique limitations. Variation in acquisition, processing and connectome metrics as well as statistical methods may impact the extent of between-subjects variability observed. For instance, for SC, the diffusion method, tractography (Bonilha et al., 2015), SC metric (Buchanan, Pernet, Gorgolewski, Storkey, & Bastin, 2014), or ROI size (Bonilha et al., 2015), have been shown to affect variability and reproducibility of SCs. FC variability across subjects is affected by the choice of metric (Marrelec et al., 2016). For example, the amount of common variance may be slightly higher when using correlation compared to mutual information for the calculation of FC. On the other hand, the common component of FC that is invariant across subjects was comparable for dynamic and static FC (Marrelec et al., 2016). Nonetheless, the correlation between SC and FC may be limited by the dynamic fluctuation of FC on short time windows (Allen et al., 2014; Deco, Kringelbach, Jirsa, & Ritter, 2016; Hutchison et al., 2013). SCs may better correlate with temporally stable rsFC (Honey et al., 2009). To this end, we considered only SC present connections in a secondary analysis, as these have been shown to have more stable resting-state FC (Shen et al., 2015).

One important question is whether increased between-subjects variation in the FC is a consequence of non-neural influences such as vascular variability or head motion (Geerligs, Tsvetanov, Cam, & Henson, 2017) or reflects real, meaningful variability in neural activation. If meaningless between-subjects variability in FC can be reduced, FC has the best chance to be able to capture subtle individual differences in SC. In addition to the corrections described in the methods, FC between-subjects variability was minimized via a secondary global signal regression (GSR) analysis (Berlin dataset, NKI Rockland dataset). Yet, lack of SC-FC subject-specific correlation in five of the six persists despite these secondary analyses.

### Future directions

Computational models that investigate how SC gives rise to FC may be particularly helpful for furthering our understanding of how individual SC and FC are linked (Jirsa, Sporns, Breakspear, Deco, & McIntosh, 2010; Kringelbach, McIntosh, Ritter, Jirsa, & Deco, 2015; Kunze, Hunold, Haueisen, Jirsa, & Spiegler, 2016; Ritter et al., 2013; Roy et al., 2014). The mechanisms by which individual FC comes about from individual SC may be the key to understanding subject-specific differences. To this end, parameters from generative models combining individual empirical SC and FC can be used (Schirner, McIntosh, Jirsa, Deco, & Ritter, 2018). Variability in these parameters have already been shown to be useful for revealing individual differences relevant for cognition (Falcon et al., 2016; Falcon et al., 2015; J. Zimmermann et al., 2018). These parameters may even exceed the predictive capacity of individual connectomes (Zimmermann et al., 2018).

### Summary

We present evidence that, in most standard datasets, the subject variation in SC may be too weak to be reflected in the FC variability. However, subject-specificity of SC-FC can be captured via fine, multi-modally parcellated data, due to greater SC variability across subjects. Nonetheless, SC and FC each show a large component that is common across subjects, which sets limitations on the extent of SC-FC subject-specificity. Implications of these findings for personalized medicine should be considered. Namely, attention to the quality of processing and parcellation methods is critical for furthering our understanding of the relationship between individual SC and FC.

## Acknowledgments

Data were provided [in part] by the Human Connectome Project, WU-Minn Consortium (Principal Investigators: David Van Essen and Kamil Ugurbil; 1U54MH091657) funded by the 16 NIH Institutes and Centers that support the NIH Blueprint for Neuroscience Research; and by the McDonnell Center for Systems Neuroscience at Washington University.

The authors gratefully acknowledge the computing time granted by the John von Neumann Institute for Computing (NIC) and provided on the supercomputer JURECA at Jülich Supercomputing Centre (JSC) (www.fz-juelich.de, Grant NIC#8344 & NIC#10276 to P.R.). The authors acknowledge the support of the NSERC grant (RGPIN-2017-06793) to A.R.M., and funding granted by the German Ministry of Education and Research (US-German Collaboration in Computational Neuroscience 01GQ1504A, Bernstein Focus State Dependencies of Learning 01GQ0971-5), the European Union Horizon2020 (ERC Consolidator Grant BrainModes 683049), Stiftung Charité/Private Exzellenzinitiative Johanna Quandt and Berlin Instititute of Health (BIH Johanna Quandt Professorship for Brain Simulation) to P.R.

